# Differences in unity: the go/no-go and stop signal tasks rely on different inhibitory mechanisms

**DOI:** 10.1101/705079

**Authors:** Liisa Raud, René Westerhausen, Niamh Dooley, René J. Huster

## Abstract

Response inhibition refers to the suppression of prepared or initiated actions. Typically, the go/no-go task (GNGT) or the stop signal task (SST) are used interchangeably to capture individual differences in response inhibition. Yet, there is some controversy if these tasks assess similar inhibitory processes. We extracted the time-courses of sensory, motor, attentional, and cognitive control networks by group independent component (G-ICA) analysis of electroencephalography (EEG) data from both tasks. Additionally, electromyography (EMG) from the responding effector muscles was recorded to detect the timing of response inhibition. The results indicated that inhibitory performance in the GNGT may be comparable to response selection mechanisms, reaching peripheral muscles at around 316 ms. In contrast, inhibitory performance in the SST is achieved via biasing of the sensory-motor system in preparation for stopping, followed by fast sensory, motor and frontal integration during outright stopping. Inhibition can be detected at the peripheral level at 140 ms after stop stimulus presentation. The GNGT and the SST therefore seem to recruit widely different neural dynamics, implying that the interchangeable use of superficially similar inhibition tasks in both basic and clinical research is unwarranted.

## 1. Introduction

Response inhibition is fundamental for purposeful behavior, enabling us to adapt rapidly to changes in the environment. By a strict definition, response inhibition refers to the suppression of a prepared or initiated action. In its broader context, inhibitory control relies on a cascade of neural processes that ultimately result in the suppression of behavior, including signal detection and discrimination, response preparation, interference control, and, eventually, response inhibition. It is likely then, that successful inhibition depends on the fine-tuned interaction of multiple control systems, depending on the specific task requirements or strategies. Yet, the majority of the scientific literature as well as neuropsychological tests treat the plethora of available inhibition tasks as interchangeable, thus inherently presuming that response inhibition is a unitary construct.

The most common behavioral paradigms to measure response inhibition are the go/no-go task (GNGT) and the stop signal task (SST). In both paradigms, the primary task is either a simple or a choice reaction task. In the GNGT, a proportion of the stimuli are replaced with a no-go stimulus, while in the SST, the go stimulus is always shown first, but may then be followed by a stop stimulus after a short stop signal delay (SSD, typically around 250-350 ms). Both the no-go and the stop stimulus instruct the participant not to respond, despite the prepared or possibly initiated go-response. The single difference between the two tasks is thus the timing of the inhibition signal relative to the go signal (0 and ~300 ms for the GNGT and SST, respectively).

The GNGT and SST are the primary tasks for measuring response inhibition and they are typically used indiscriminately. However, there is some controversy whether they activate the same or different inhibitory control mechanisms. A distinction has been proposed with the GNGT capturing action restraint, whereas the SST supposedly relies on action cancellation (Schachar et al., 2007). Action restraint in the GNGT refers to a decision of whether or not to respond. In contrast, there is no such decision in the SST as the default decision is always to respond, so action cancellation refers to the suppression of this already initiated response. Behavioral evidence supports the dissociation of these mechanisms, as performance in one task can deteriorate without apparent deficits in the other (Krämer et al., 2013; Littman and Takács, 2017). Furthermore, pharmacological manipulations suggest that action restraint relies on serotonergic and action cancellation on noradrenergic neurotransmitter signaling (Eagle et al., 2008). However, human functional neuroimaging studies indicate a functional convergence of neural processing within a single inhibitory control network. Studies directly comparing the two tasks report common activations in inferior frontal cortex (IFC) and/or insula, middle frontal cortex, (pre-) supplementary motor area (pre-SMA), cingulate cortex, dorsolateral prefrontal cortex, and posterior parietal regions (Cai et al., 2014; Dambacher et al., 2014; McNab et al., 2008; Nee et al., 2007; Rubia et al., 2001; Sebastian et al., 2013; Swick et al., 2011; Zheng et al., 2008). Separate networks for either task have been reported as well, although the results across different studies tend to be less convergent. Rubia et al. (2001), for example, report left-hemispheric dominance for no-go trials over stop trials, while Swick et al. (2011) stress the role of right medial frontal and right inferior parietal areas for no-go trials. Stronger activity for stop than for no-go trials has been reported in right insula and striatum (Sebastian et al., 2013), left insula (Swick et al., 2011), cingulate (Dambacher et al., 2014), thalamus (Dambacher et al., 2014; Swick et al., 2011), subthalamic nucleus, and substantia nigra (Guo et al., 2018). These observations lead to two competing interpretations: given the prominence of the IFC-preSMA network in response inhibition (Aron et al., 2014, 2004), it is likely that a common action inhibition network is activated in both tasks and the variations in brain activation reflect differences in general task demand or difficulty. In contrast, task-specific activations beyond the shared networks may suggest that the tasks recruit different mechanisms, but the common activations reflect shared task aspects, such as stimulus processing, attentional capture, or response evaluation.

It seems that electroencephalography (EEG) is better suited than fMRI for resolving the train of cognitive processes at a fine temporal scale. A majority of response inhibition findings has focused on the event related potentials (ERPs) N2 and P3 that are elicited reliably in both the GNGT and the SST (Huster et al., 2013). The P3 onset, particularly, has been proposed as the marker for inhibition (Wessel and Aron, 2015). When the two tasks are compared directly, the P3 amplitude tends to be larger in the SST than in the GNGT (Cunillera et al., 2016; Enriquez-Geppert et al., 2010; Johnstone et al., 2007). Further, Wessel (2017) manipulated the task parameters of the GNGT and found that the stop-related P3 was elicited in the GNGT only if the probability of no-go stimuli was low and the task pace was fast enough to evoke a prepotent response tendency. However, others have argued that the P3 appears too late to index a genuine inhibitory process (Filipović et al., 2000; Huster et al., 2019; Raud and Huster, 2017; Skippen et al., 2019). While the P3 onset is typically detected at around 200-300 ms, corticomotor excitability is reduced already at around 150 ms inhibitory signal onset (Coxon et al., 2006; Fujiyama et al., 2011; Hoshiyama et al., 1997, 1996; Macdonald et al., 2014; van Campen et al., 2013; van den Wildenberg et al., 2010; Yamanaka et al., 2002). This coincides with the estimation of inhibition latencies of around 150 ms from partial response electromyography (prEMG) indices (Raud and Huster, 2017). PrEMG refers to EMG bursts in trials where no overt button press is registered. These have been observed in a number of conflict tasks and are often considered erroneous response activations that are inhibited before they fully develop into an error (hence the previously used term ‘partial error’; Burle et al., 2002; Hasbroucq et al., 1999; Rochet et al., 2014). In the SST, however, these bursts indicate correct response activation in reaction to the go signal that then gets cancelled in response to the stop signal (Atsma et al., 2018; De Jong et al., 1990; Raud and Huster, 2017). Correspondingly, while the onset latency of such EMG reflects response initiation, the starting point for its decline (quantified as the prEMG peak latency) may indicate the time-point of inhibitory effects in the periphery.

The focus on the N2 and P3 in the response inhibition literature may stem from the fact that these components are relatively large in amplitude and thus tend to dominate over other EEG markers of different stages of cognitive processing. Indeed, studies that attempt to de-mix the various sources of EEG activity report a much richer set of activations, particularly in early time-windows just after the presentation of the inhibitory signal (Albares et al., 2014; Huster et al., 2017, 2014). Additionally, recent studies highlight the importance of other ongoing processes for successful inhibition, such as an attentional bias towards fast stimulus detection (Langford et al., 2016a, 2016b; Skippen et al., 2019) or motor preparation processes (Liebrand et al., 2018).

Here, we present time-resolved neural network activity profiles for both the GNGT and the SST that represent different stages of cognitive processing, spanning from perception to action and to higher cognitive control functions, including response inhibition. Time-variant functional networks were extracted by group independent component analysis (G-ICA; Eichele et al., 2011; Huster et al., 2015; Huster and Raud, 2018) of the combined EEG activity from both tasks, and the resulting independent component (IC) time-courses were compared statistically by permutation testing. Distributed source analysis was performed on the topographical maps of the ICs to aid their interpretability as functional networks. In addition, EMG was recorded from the responding hands. By comparing the temporal profiles of the ICs and prEMG activity within and between the two tasks, we aimed to differentiate if these tasks rely on shared or different inhibition mechanisms. A shared inhibition mechanism would be reflected in similar temporal dynamics of no-go and stop activity before or at the time of the inhibition latency. In contrast, the GNGT and SST may rely on different mechanisms, namely action restraint and cancellation, respectively. This would result in largely different IC timecourses when no-go and stop trials are contrasted directly. Note that under a strict interpretation of this alternative, processes other than inhibition are expected to show comparable time-courses between the tasks, while a more lenient interpretation would also expect other processes to be recruited in a diverging manner.

## 2. Methods

### 2.1 Sample

Data was collected from 37 participants (20 females, 17 males) between the age of 19 and 35 years (average age 26.8). The SST data is also used in Huster et al. (2019). All participants were right-handed, reported no psychiatric or neurological disorders and had normal or corrected-to-normal vision. Data from four participants was discarded due to low performance on the stop signal task (go accuracies < 80%) and one participant was excluded as he did not complete the full procedure. The final sample thus consists of 32 participants. The participants came for two sessions within one week, one for EEG and one for MRI. Prior to the experiments, all participants gave written informed consent. Participants received monetary compensation for study participation. The study was conducted in accordance with the ethical standards of the Declaration of Helsinki and was approved by the internal review board of the University of Oslo.

### 2.2 Tasks and procedure

All participants performed both the GNGT and the SST in a single session, with the order of tasks counterbalanced across participants. Each task lasted for about 30 minutes. Task administration was controlled using E-prime 2.0 (Psychology Software Tools, Pittsburgh, PA).

The go-stimuli were green arrows, pointing either to the left or to the right (arrowhead 3 x 3.5 cm and base 3 x 1.5 cm). The no-go/stop stimuli were blue arrows of the same size as the go stimuli. The participants sat at a viewing distance of approximately 80 cm from the screen. All stimuli were presented in the center of the screen and a fixation cross was presented at all other times during the experiment.

In the GNGT, participants were instructed to respond as quickly as possible via a button press with the thumb of the hand corresponding to the direction of the arrowhead. In some trials, a no-go stimulus appeared instead of a go stimulus, signaling the participant not to respond. In the SST, the primary task was the same as in the GNGT, namely to react to the go stimuli with left or right button presses. In stop trials, the stop stimulus appeared after the go stimulus after a short delay (stop-signal delay; SSD), instructing the participant to suppress their initiated response.

In both tasks, all stimuli were presented for 100 ms and the inter-trial interval was randomly set to a value between 1500-2500 ms. In the SST, the SSD was varied according to a performance tracking procedure that would result in a stopping success of 50%. The initial SSD was set to 250 ms and was increased or decreased in steps of 50 ms if stopping in the preceding stop trial was successful or unsuccessful, respectively. The minimum possible SSD was the go stimulus duration (100 ms) and the maximum was set to 800 ms. The SSD tracking was done separately for stop left and stop right trials.

Each task had 800 trials in total, with 600 go trials and 200 no-go or stop trials (25% of all trials). There was an equal number of left and right hand trials with 300 go left/right and 100 no-go or stop left/right trials. The trials were equally distributed over 10 blocks (80 trials per block) and participants received feedback after each block, instructing them to be faster if the average go reaction time of the previous block exceeded 500 ms. In addition, instantaneous feedback ‘Too slow!’ was presented after go omissions or if the RT of a given trial exceeded 1000 ms. Pauses of self-regulated duration were introduced after each block. A longer break with a duration of 5-10 minutes was allowed in-between the two tasks. Prior to both tasks, participants completed a short training session of 20 trials to introduce them to the tasks. In the SST, it was additionally stressed that it was not possible to be correct all the time and that it is important to be both fast and accurate, to prevent excessive waiting for the stop stimulus.

### 2.3 Data acquisition

EEG and EMG were recorded using a Neuroscan SynAmps2 (Compumedics) amplifier. The data was digitized at 2500 Hz. EEG was measured from 64 Ag/Ag-Cl electrodes, positioned according to the extended 10-20 system with two horizontal EOG channels placed beside the left and the right eye. All EEG electrodes were referenced online against an electrode placed at the nose-tip and their impedances were kept below 5 kOhm. Electrode AFz served as the ground electrode.

For EMG recordings, bipolar Ag/Ag-Cl electrodes were placed on the skin surface above the *abductor pollicis brevis*, parallel to the belly of the muscle. The ground electrode was placed on the left arm. Support was provided for the participants’ arms to reduce spurious baseline muscle tension.

Structural MR images were acquired on a Philips Ingenia 3T scanner with a 32-channel Philips SENSE head coil. The T1-weighted image was obtained with a sequence of 184 sagittal slices of 1mm thickness, and an in-plane resolution of 256 x 256 at a FoV of 256mm, which resulted in a voxel size of 1×1×1 mm (TE 2.2 ms, TR 4.66 ms, flip angle 8°).

### 2.4 Data analysis

#### 2.4.1 Behavior

The following behavioral measures were extracted: go trial accuracies, correct go trial reaction times (RT), no-go accuracies, stop accuracies, and in case of the SST, unsuccessful stop trial RTs and the SSRTs. SSRTs for each subject were estimated separately for left and right hand responses using the integration method. That is, the mean SSD was subtracted from the go RT distribution percentile corresponding to the probability of unsuccessful stopping (Band et al., 2003; Logan and Cowan, 1984). According to an initial analysis, right hand responses showed slight advantage compared to the left hand responses in terms of higher go accuracies and faster RTs. This effect was small, however (in the order of 2% and 10 ms for accuracies and RTs, respectively) and similar in both tasks. Thus, all behavioral variables are reported as an average over left and right hand trials to reduce redundancies. Differences in go accuracies and RTs were compared between the tasks using paired t-tests.

#### 2.4.2 EMG analysis

EMG channels were filtered between 10-200 Hz and resampled at 500 Hz. Data was first segmented relative to go or no-go stimulus onset (+/-2.5 s). Trials with amplifier saturation were discarded from the analysis (1.61% of all trials in the GNGT and 1.78% in the SST). A moving average procedure was applied where, for each time point, the average root mean square over +/-5 neighboring points was calculated. The resulting waveforms were then divided by the trial-specific average of pre-go (or nogo) activity from −200 to 0 ms, and then z-scored across all trials and time-points, separately for each hand. Data was then re-segmented into conditions with a time-window of −200 to 1000 ms relative to go-stimulus in go and no-go trials and −600 to 1000 ms relative to stop-stimulus in stop trials. The trials were baseline corrected by subtracting the average trial-specific baseline of −200-0 ms in go/no-go trials and −600 to −400 ms in stop trials. The different baseline windows were selected so that all trials would be baseline corrected to pre-go (or no-go) activity, which in stop trials was the stop onset minus the average SSD of 400 ms. An automatic algorithm was used to detect EMG bursts, regardless of the trial type and given response. An EMG response was identified when the z-scored and baseline corrected activity exceeded the threshold of 1.2. This threshold was chosen based on a visual inspection of the data, and random trials were checked manually to confirm the algorithm’s performance. If an EMG burst was detected in a given trial, two variables were extracted: EMG onset latency (the first point over the threshold) and EMG peak latency. Additionally, EMG burst frequency was defined as the percentage of trials with an EMG burst relative to the total number of trials in a given condition. Trials where EMG onset started earlier than the go or no-go stimulus and where prEMG peak was earlier than the stop stimulus were discarded from further analysis.

For completeness, EMG frequencies were estimated for all trials (go, no-go, stop) and both for selected (e.g. left hand in go/no-go/stop left trials) and unselected hand (e.g. right hand in go/nogo/stop left trials, Table 2). The EMG frequencies in selected hand go trials were estimated to validate the performance of the automatic EMG algorithm. The prEMG frequency in the unselected hand for all trials was estimated to investigate the possibility of erroneous activations during the response selection stage. EMG indices were averaged over left and right hand trials (since an initial analysis showed no differences between them).

Statistical analysis was performed on the selected hand prEMG in the no-go/stop trials and in the unselected hand prEMG in the go trials, to account for both, correctly inhibited prEMG responses in the no-go/stop trials and erroneous prEMG in the go trials due to response competition. This resulted in a full repeated-measures ANOVA with factors TASK (GNGT, SST) and TRIAL (go unselected hand, no-go/stop selected hand) for the detection frequencies. An additional analysis was performed on the prEMG peak latencies; however, the go unselected hand condition in the SST was discarded due to the low number of trials that would yield unreliable latency estimates. This resulted in a one-way ANOVA with a hybrid factor of TASK-TRIAL (GNGT-go-unselected, GNGT-no-go-selected, SST-stop-selected). The ANOVAS were calculated with the package *ez* in the R Project for Statistical Computing.

#### 2.4.3 EEG preprocessing

EEGLAB (version 14.1.2) was used for data preprocessing (Delorme and Makeig, 2004). EEG channels were filtered between 0.1 to 80 Hz, resampled at 500 Hz and re-referenced to the common average of all EEG channels. Data was first epoched from −2.5 to 3.5 s around all go stimuli in the SST, and around all go and no-go stimuli in the GNGT. ICA was run on the segmented data and components capturing eye or muscle artifacts, as identified by visual inspection, were rejected. The remaining components were back-projected to the channel domain. Data was then re-segmented with a timewindow of −0.2 to 1 s relative to go stimulus onset in go trials and no-go/stop signal onset in the no-go/stop trials. Baseline correction was applied by subtracting the mean pre-stimulus activity from each trial. Trials with an absolute value exceeding the threshold value of 75V were discarded as artifacts (0.5% GNGT, 1.1% SST).

#### 2.4.4 G-ICA

G-ICA was performed using publicly available code for Matlab (Huster and Raud, 2018). For the G-ICA, it is necessary to have an equal number of trials for all participants in all conditions. Thus, a subselection of trials was used for G-ICA, with 44 trials for each of the two tasks (the lowest number of artefact-free trials available across all participants and conditions). Note that this sub-selection was only for the purpose of calculating the G-ICA solution; once computed, the back-projection of the ICs was done on each individual’s whole data set. The no-go and stop trials were selected randomly; however, only the go trials with the reaction times closest to the group average were selected for the G-ICA. This is because G-ICA tends to perform poorly on the activity that is not homogeneously time-locked to the stimulus onset, such as reaction times (Huster et al., 2015). G-ICA estimates a common component structure across subjects by combining subject-specific and group-level principal component analysis (PCA) with group-level ICA. Ten components were extracted from the single-subject PCA as this explained, on average, 90% of each individual’s data. To determine the number of group components, we used the ICASSO procedure (Himberg and Hyvarinen, 2003) where we extracted a set of components from 5 to 15, 100 times each. For each number of components, the stability over the 100 runs was checked by the means of agglomerative clustering with average linkage. The number of components that resulted in the most stable cluster solution (determined by the visualization of the clusters and the estimate of the stability index Iq) was then selected for the G-ICA as used for the final analyses. As a result of this procedure, 11 components were extracted.

Although G-ICA performs reasonably well in dissociating EEG components with temporal overlap (Eichele et al., 2011; Huster et al., 2015), there still remained the possibility that due to the fast sequential presentation of go and stop trials in the SST, the stop trial data may be contaminated by systematic go-related EEG activity. Thus, to further separate the go and stop processes, a regression analysis was performed instead of the standard trial averaging to obtain condition-specific component regression ERPs (rERP). First, individual IC time-courses were computed from the continuous EEG data of each participant by applying the participant-specific decomposition matrices obtained via G-ICA. Then, automatic rejection of artifacts on the continuous data was performed using the EEGLAB *pop_rejcont* function by rejecting data-points where the power of higher frequencies (20-80 Hz) exceeded 15 dB (~10% of all data, including breaks between the blocks). Finally, beta weights for each condition were estimated by regression using the EEGLAB plugin rERP (Smith and Kutas, 2015). The weights were estimated separately for left and right hand trials, as hemispheric lateralization effects may be captured by separate ICs. The beta weights were estimated for each event of interest from −200 to 1000 ms. To prevent overfitting due to an over-defined model, regularized regression with L2-norm penalization was used. The regularization parameter lambda was determined as the one yielding the highest R^2^ in a grid-search with 5-fold cross-validation.

#### 2.4.5 IC statistical analysis

Each component’s condition-specific time-course from −200 to 600 ms was tested by permutation tests combined with threshold-free cluster enhancement (tfce). Tfce takes the autocorrelation of the component data into account and enhances the signal at each time-point depending on the activity of the neighboring data-points (Smith and Nichols, 2009). A statistical test is then performed on each data-point separately and a correction for multiple comparisons is applied to the resulting p-values. In contrast to the cluster-mass inference procedure, inference is done on each data-point (as opposed to the whole cluster), which has the advantage that the first or last time-point of the cluster of significant activations is meaningful and can be interpreted as the onset or offset of condition differences, respectively. The analysis was performed using the PALM software (Winkler et al., 2014). In the GLM, we tested the effect of the factors TRIAL (go vs stop/no-go), SIDE (left vs right) and the interaction of TRIAL x SIDE separately for the GNGT and SST. The parameters for tfce were H = 2, E = 1, and the rERP time-courses were permuted 10000 times within participants. The resulting two-tailed p-values were corrected for multiple comparisons using a false discovery rate (fdr) correction at an alpha value of 0.05. We only report clusters with significant effects that span more than 20 ms. Next, a conjunction analysis for the GNGT and SST was performed in order to identify time-intervals where no-go activity was different from go activity in the GNGT *and* stop activity was different from go activity in the SST. That is, the t- and p-values from the go vs. no-go and go vs. stop comparison were combined by taking the minimum absolute t-value and corresponding p-value per time point (Nichols et al., 2005). Note that the absolute t-value is blind to the direction of the effect and may falsely identify a conjunction where the main effects for two tasks were in the opposing directions. For this reason, the conjunction analysis was performed twice for each main effect, once for positive and once for negative t-values. The resulting p-values were fdr-corrected at an alph-alevel of 0.025, which corresponds to two-tailed alpha value of 0.05. Lastly, another model with factors TRIAL (no-go/stop) and SIDE (left/right) was fit to directly test the differences between the IC time-courses of the two types of inhibition. The procedure and parameters for tfce, permutations, and multiple comparison correction were the same as before.

The results from the permutation analysis suggested pre-stimulus differences between go and stop trials in multiple ICs. To further investigate the dynamics of these ICs, two additional analyses were performed on a post-hoc basis. For the first one, a median split was computed for each participant’s go RTs and ‘slow’ and ‘fast’ time-courses were extracted separately by the rERP procedure. These were then subjected to a similar permutation testing as described above, with the factors RT (slow vs fast) and SIDE (left vs right). Similarly, unsuccessful stop trial beta weights were extracted by the rERP procedure and subjected to statistical testing with factors STOP (successful vs unsuccessful) and SIDE (left vs right).

#### 2.4.6 IC source localization

Each participant’s cortical surface was extracted automatically from the T1 MRI image using the Freesurfer analysis suite (http://surfer.nmr.mgh.harvard.edu/), and the individual anatomies were imported to Brainstorm for source analysis (Tadel et al. 2011; http://neuroimage.usc.edu/brainstorm). The standard EEG electrode locations were co-registered with each participant’s head surface and boundary element headmodels were constructed with OpenMEEG (three layers with 1922 vertices each; relative conductivities: scalp = 1, skull = 0.0125, brain = 1; Gramfort et al., 2010; Kybic et al., 2005). Source estimation was done for 15002 dipoles distributed along the cortical surface using the individual IC time-courses that were back-projected to the channel domain and averaged across trials. Since each IC is characterized by a unique topography that is constant over time and conditions, we selected the condition and the time-window for each subject at which the IC’s signal-to-noise-ratio was the highest. This was calculated by dividing the IC time-course into bins of 50 ms and dividing the mean absolute value of a given bin by the standard deviation of the baseline period. The sources were estimated for the resulting condition using sLORETA with unconstrained source orientations. Diagonal noise-covariance matrix was used with the regularization parameter set to 0.1. The resulting 3-dimensional dipole values were projected to a single value using singular value decomposition and averaged over time-points selected time-bin. The source maps were then projected to a standard anatomical template (ICBM152) and down-sampled to the Destrieux atlas for anatomical labeling and interpretation. As distributed source modeling algorithms tend to falsely identify strong responses at the edges of the forward model, such activity patterns were considered unreliable and discarded from further analysis. For the remaining regions, t-tests for deviations from zero were performed to assess the significance of the source constellation of each IC at the group level. The resulting p-values were fdr-corrected for multiple comparisons with an alpha level of 0.05.

## 3. Results

### 3.1 Behavior

Behavioral results are listed in Table 1. Overall, participants showed good task performance as indicated by go accuracies > 94% in both tasks and an average no-go accuracy of 99%. Stop accuracies were close to 50% and reaction times in unsuccessful stop trials were faster than go reaction times for each participant and at the group level (t (31) = 21.48, p < 0.001, d = 3.80). This is in good compliance with the horse race model assumption stating that the reaction times in the unsuccessful stop trials should correspond to the left side of the go RT distribution. The average SSRT was 194 ms. While go accuracies were similar between the GNGT and the SST (t(31) = −0.90, p = 0.375, d = −0.159), RTs were considerably slower in the SST than in the GNGT (t(31) = 13.89, p < 0.001, d = 2.46).

**Table 1.**
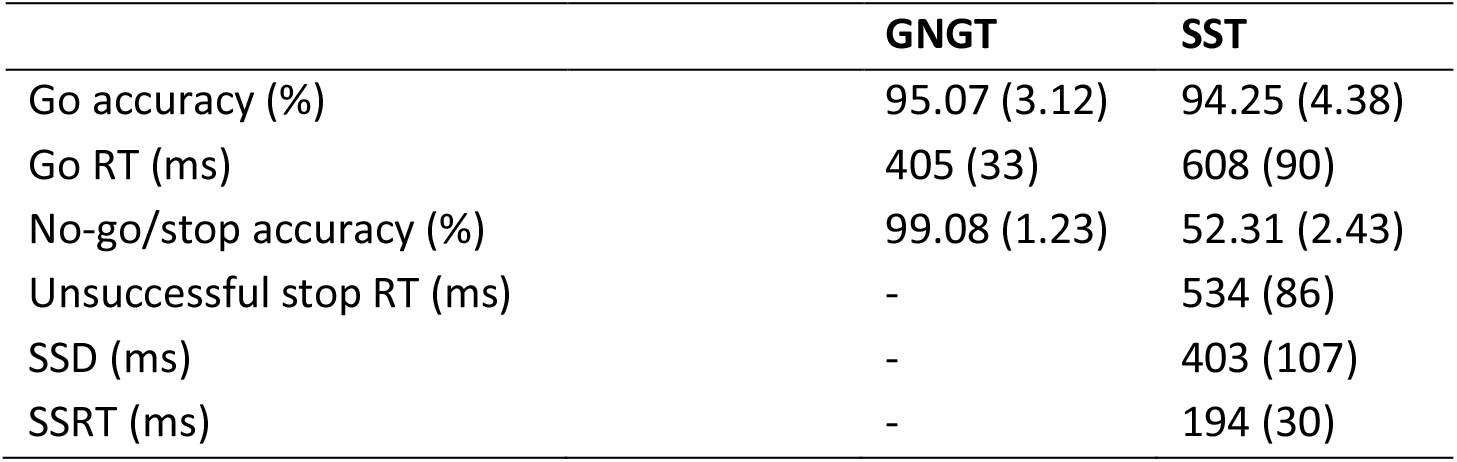
Summary statistics (means and standard deviations) of the behavioral measures.

|  | GNGT | SST |
| --- | --- | --- |
| Go accuracy (%) | 95.07 (3.12) | 94.25 (4.38) |
| Go RT (ms) | 405 (33) | 608 (90) |
| No-go/stop accuracy (%) | 99.08 (1.23) | 52.31 (2.43) |
| Unsuccessful stop RT (ms) | - | 534 (86) |
| SSD (ms) | - | 403 (107) |
| SSRT (ms) | - | 194 (30) |

### 3.2 EMG

#### 3.2.1 Overt response EMG

All extracted EMG measures are listed in Table 2 and EMG time-courses are depicted in Figure 1. The visualization of the time-courses of the selected hand go trials corroborate the behavioral findings of later responses in the SST than in the GNGT. Go reaction times in the SST also appear to have larger variability that contributes to the reduced amplitudes of the averaged EMG wave-form. The automatic EMG detection algorithm performed well, as indicated by the detection of EMG bursts in around 97% of correct go trials. The below 100% detection rate implies a slightly conservative performance of the algorithm; misses were due to random noise or elevated muscle activity in the pre-go baseline period. Baseline RMS values were similar for the GNGT and SST (mean baseline RMS across all trials were 9.84 and 9.66 for the GNGT and SST, respectively).

**Figure 1.**
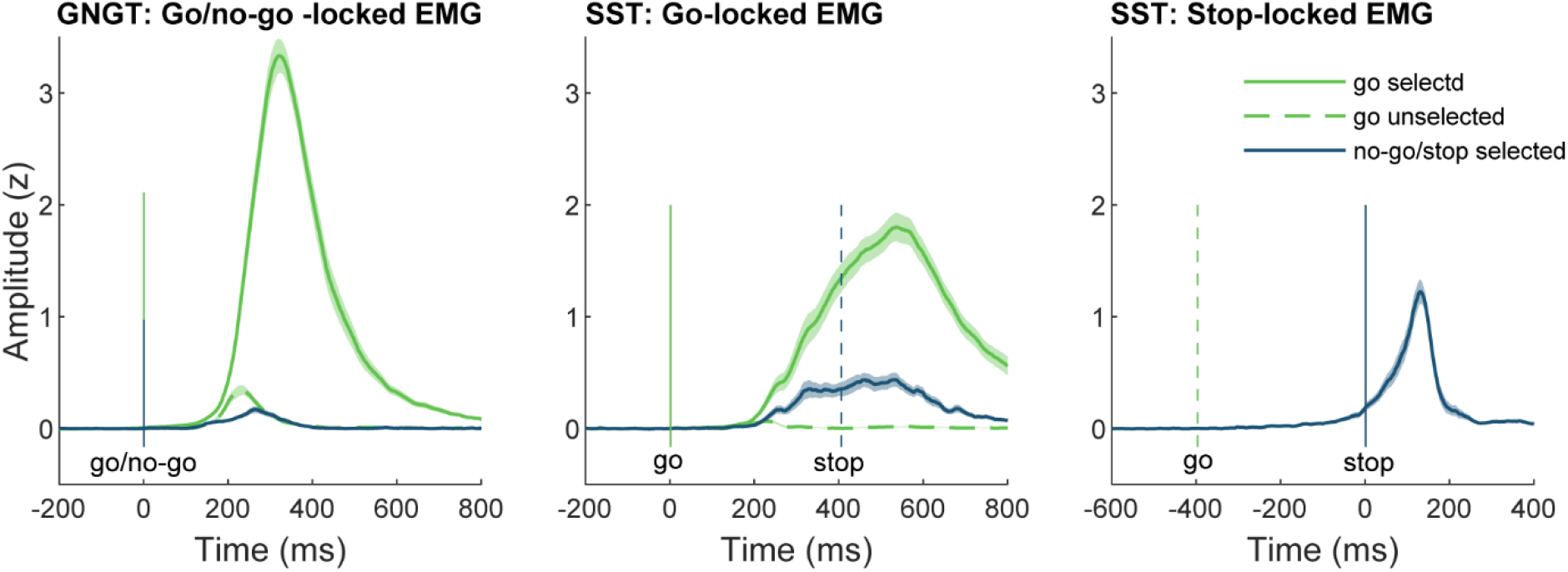
Averaged EMG time-courses. The shaded areas represent standard errors. The vertical solid lines represent the time-locking event and the vertical dashed lines represent the average SSD relative to the time-locking event.

**Table 2.** Mean values and the standard deviations of the EMG measures. Onset latencies are calculated relative to the go or no/go stimulus. Peak latencies are calculated relative to the event of interest, i.e relative to go stimulus in go trials, no-go stimulus in no-go trials, and stop stimulus in stop trials. N.e. -; not estimated due to low number of trials. Values in bold are trials where an overt response was given; the other values represent prEMG activity without an overt response.

|  | GNGT |  | SST |  |
| --- | --- | --- | --- | --- |
|  | Go | No-go | Go | Stop |
| **Selected hand** |  |  |  |  |
| Frequency (%) | **97.55 (3.38)** | 14.18 (10.43) | **96.55 (3.89)** | 44.57 (18.68) |
| Onset (ms) | **282 (36)** | 285 (78) | **452 (92)** | 455 (86) |
| Peak (ms) | **350 (40)** | 316 (88) | **545 (91)** | 140 (29) |
| **Unselected hand** |  |  |  |  |
| Frequency (%) | 20.75 (12.62) | 6.65 (7.55) | 7.75 (9.16) | 4.82 (7.44) |
| Onset (ms) | 258 (47) | n.e. | n.e. | n.e. |
| Peak (ms) | 289 (56) | n.e. | n.e. | n.e. |

#### 3.2.2 PrEMG

The primary measures of interest were the prEMG bursts, that is, EMG bursts in trials without registered button presses. Such prEMG activity was clearly visible in the averaged EMG time-courses and they were especially pronounced in the SST when the time-courses were locked to the stop signal onset (Figure 1). PrEMG was more frequent in successful stop trials (45%) than in successful no-go trials (14%). Interestingly, prEMG responses were detected also in the unselected hands in go trials (21% in GNGT, 8% in SST), reflecting response competition in go trials. The ANOVA on the prEMG frequencies found a main effect of the TASK (F(1,31) = 17.42, p < 0.001, 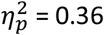) and TRIAL (F(1, 31) = 131.44, p < 0.001, 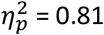), as well as the interaction of the two (F(1,31) = 190.32, p < 0.001, 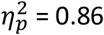). Post-hoc tests confirmed that unselected hand prEMG in go trials was more frequent in the GNGT than the SST, while selected hand prEMG occurred more often in the stop than in the no-go trials (all p’s < 0.005).

PrEMG peak latencies were also different between the trials. For this analysis, the unselected hand go trials in the SST were excluded due to the low detection rates that would yield unreliable latency estimates, resulting in a one-way ANOVA with a hybrid TASK-TRIAL factor (GNGT-go unselected, GNGT-no-go selected, SST-stop selected). This analysis revealed a main effect of this 3-level factor on prEMG peak latencies (F(2,62) = 123.12, p < 0.001, 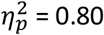). Post-hoc comparisons confirmed that the prEMG responses peaked at the earliest in stop trials (140 ms relative to stop), then in GNGT go trials (unselected hand; 289 m–s relative to go) and then in no-go trials (316 ms relative to no-go; all p’s < 0.006).

In sum, partial responses of the inhibited hand were detected both in the no-go and stop trials. This EMG activity peaked earlier in the SST than in the GNGT, perhaps indicating fast response inhibition in the SST. Furthermore, the unselected hand prEMG responses indicate stronger response competition in go trials of the GNGT than the SST.

### 3.3 EEG

#### 3.3.1 G-ICA descriptions

Both, no-go and stop trials elicited the typical N2/P3 complex that was strongest at central midline electrodes in the standard ERPs (Figure 2). Notably though, go trials in the GNGT also elicited a relatively prominent P3 peak, although its topography was more parietally focused compared to the no-go P3. Eleven independent components (ICs) were extracted via G-ICA. The relevant IC time-courses, topographies, and source constellations are depicted in Figure 3. Based on the time-courses, topographies, and sources, the resulting ICs can be interpreted as sensory (IC1), motor control (IC2 and IC3), and attentional or cognitive control components (IC4-IC7). The remaining four components were relatively small in amplitude, had rather unspecific topographies, and showed no or only minor differences between tasks and/or conditions. Thus, although these ICs showed some stability across the different runs of ICASSO, they appeared to have only minor contributions to the general task processing. Alternatively, they may represent mixed processes due to very different contributions to the group IC from individual participants, hampering their interpretability as functional networks. Given the low contribution of these ICs and their ambiguous interpretation, these four components will not be discussed further.

**Figure 2.**
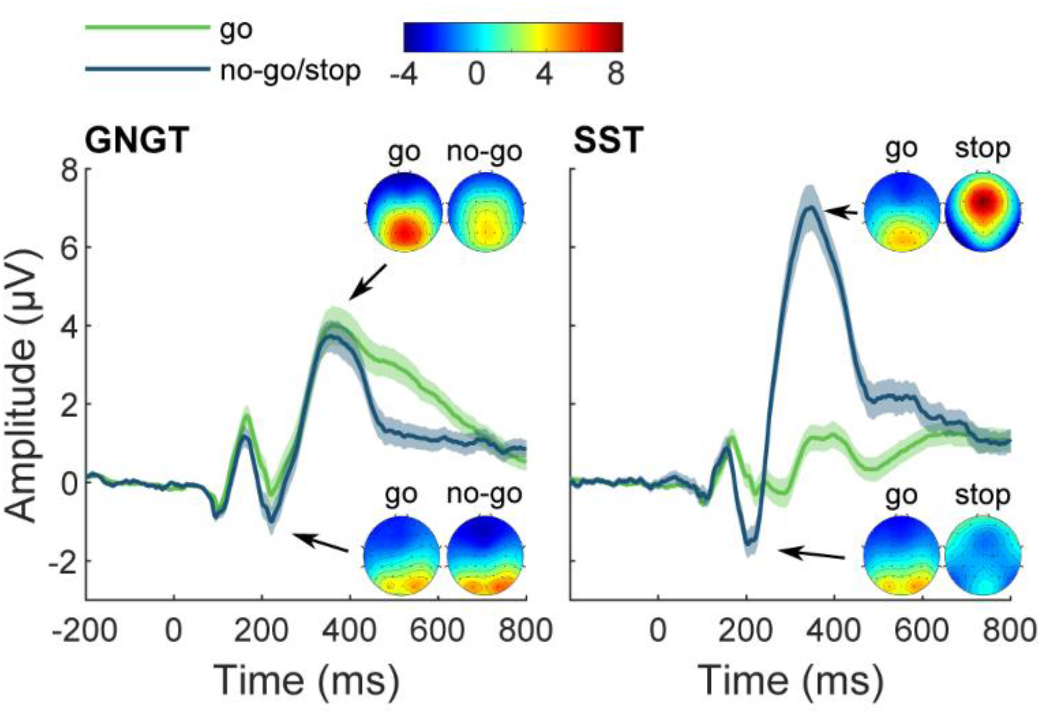
Event-related potentials (ERPs) at electrode Cz. The topographies represent the voltage distribution at all electrodes at the peak latencies of the N2 and P3 (as indicated by the arrows).

**Figure 3.**
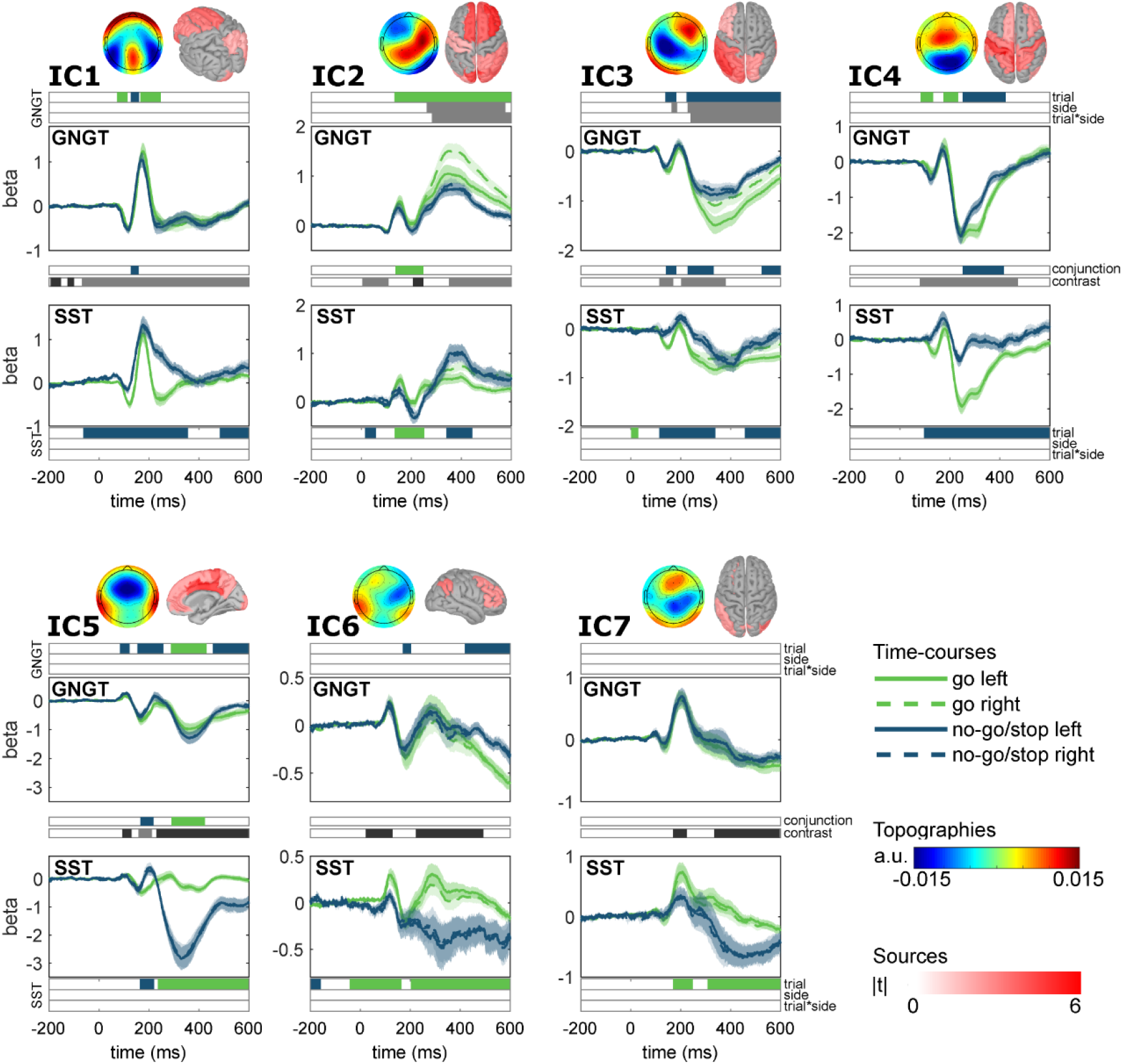
The rERP time-courses, topographies (averaged IC weight matrices) and source constellations of the group-ICs. The bars above the GNGT and below the SST time-courses represent the permutation testing results for the model TRIAL + SIDE + TRIAL*SIDE where the colors indicate significant time-intervals for each effect. The two bars in between the time-course plots of each component represent the main effect of TRIAL in the conjunction (go vs no-go AND go vs stop) and the contrast (no-go vs stop) analysis. Green represents intervals where go > no-go/stop and blue represents intervals where stop > go. In the contrast analysis, black represents intervals where no-go > stop, and gray represents intervals where stop > no-go. The source localization results are masked so that only significant regions are marked in red.

In the following, we first describe each IC’s topography, source constellation, and a global time course to derive a putative interpretation in terms of its functional neuroanatomy. Then, we describe the conditional effects in the GNGT and SST, as well as the conjunction of these effects across both tasks. Lastly, we describe the differences of the IC time-courses directly contrasting the no-go and stop trials. Since the sign of the IC time-courses are somewhat arbitrary (the true deflection on the scalp corresponds to the multiplication of the IC time-course and the topography), we refrain from describing the peaks of the time-courses as positive or negative

##### IC1: Early sensory component

IC1’s time-course and topography were reminiscent of the typical early sensory ERP complex (P1 and N1) with strongest activation in the occipital and parietal electrodes. The source localization results indicated sources in frontal (bilateral frontal-superior gyrus, bilateral anterior cingulate, bilateral insula, left middle- and inferior-frontal cortex) and parieto-occipital areas (bilateral posterior cingulate, precuneus, occipital cortex and right superior parietal cortex).

**Table 3.** The results of permutation testing and source localization. For each IC, the region with the maximum t-value, it’s corresponding anatomical label (corresponding to the Destrieux atlas) and Broadmann area (BA) are reported. The permutation testing results indicate the time-periods (ms) of statistical differences of the model TRIAL (go vs no-go/stop) + SIDE (left/right) + TRIAL*SIDE. The conjunction refers to significant differences between go vs no-go AND go vs stop, and the contrast refers to the significant differences between no-go and stop trials. As no effects for the SIDE or TRIAL*SIDE occurred in the SST, conjunction and contrast model, these columns are omitted from the table. Alpha_fdr_ = 0.05.

| IC | max t | label | BA | GNGT<br>trial | GNGT<br>side | GNGT<br>trial x side | SST<br>trial | CONJUNCTION<br>trial | CONTRAST<br>trial |
| --- | --- | --- | --- | --- | --- | --- | --- | --- | --- |
| 1 | 4.978 | Left lateral<br>anterior fissure | 45 | 72 – 114<br>128 – 160<br>166 – 248 | - | - | -62 – 356<br>484 – 600 | 128 – 158 | -194 – -152<br>-126 – -100<br>-68 – 600 |
| 2 | 6.470 | Left anterior<br>cingulate gyrus<br>and sulcus | 32 | 132 – 600 | 262 – 576 | 282 – 600 | 14 – 58<br>134 – 250<br>340 – 444 | 136 – 248 | 4 – 108<br>206 – 248<br>350 – 600 |
| 3 | 8.443 | Left lateral<br>posterior fissure | 40 | 138 – 182<br>224 – 600 | 164 – 186<br>228 – 600 | 240 – 600 | 2 – 30<br>114 – 338<br>456 – 600 | 142 – 182<br>228 – 332<br>524 – 600 | 114 – 170<br>202 – 380 |
| 4 | 7.156 | Left postcentral<br>sulcus | 1, 5 | 82 – 132<br>174 – 232<br>250 – 422 | - | - | 96-600 | 250-416 | 80 – 472 |
| 5 | 6.556 | Left posterior<br>middle cingulate | 24 | 84 – 122<br>154 – 258<br>288 – 430<br>454 – 600 | - | - | 164 – 220<br>236 – 600 | 166 – 218<br>290 – 424 | 94 – 130<br>158 – 212<br>230 – 600 |
| 6 | 4.004 | Right parietal<br>inferior angular<br>gyrus | 39 | 170 – 202<br>418 – 600 | - | - | -198 – -158<br>-42 – 164<br>202 – 600 | - | 22 – 128<br>222 – 492 |
| 7 | 4.581 | Right subparietal<br>sulcus | 31 | - | - | - | 170 – 248<br>308 – 600 | - | 170 – 224<br>334 – 596 |

##### IC2 and IC3: Motor control components

IC2’s topography spanned diagonally over right frontal and central parietal electrodes. IC3 showed a very similar, almost symmetrical topography over the left hemisphere. The time-courses of IC2 and IC3 were also similar, but opposite in handedness, with IC2 having the strongest activity in right hand go trials and IC3 in left hand go trials, the differences being most prominent in the GNGT. This suggests that IC2 and IC3 play a role in motor control. However, the source constellation suggests a wider sensory-motor-frontal network with significant clusters for IC2 in bilateral anterior cingulate, right middle-frontal cortex, left motor and somatosensory cortex, bilateral middle-occipital cortex and cuneus. IC3 was localized to left fronto-lateral and parietal cortices, bilateral insula, as well as bilateral middle-parietal, cuneus and middle-occipital regions. In sum, these two components appear to form lateralized networks with activations in the right frontal and parieto-occipital areas (as well as left motor and somatosensory areas) for IC2, and left frontal and parieto-occipital areas in case of IC3.

##### IC4, IC5, IC6 and IC7: Attentional and cognitive control components

IC4 had a parietal topography and sources in the bilateral somatosensory cortex, bilateral superior-parietal cortex and right lateral prefrontal cortex, likely corresponding to the fronto-parietal attentional network. IC5 appeared to capture the N2/P3 complex. The source localization identified a widespread cluster that covered most of the cortical areas, with strongest activations in the middle and posterior cingulate. Lastly, there were two components, IC6 and IC7, which exhibited right frontal topographies. IC7 was located to the parietal and occipital areas despite its frontal topography. IC6, however, showed two distinct clusters in the right ventro-lateral frontal cortex and right inferior-parietal cortex and might therefore correspond to the right-lateralized stopping network.

#### 3.3.2 Task-specific IC dynamics

*GNGT*. The time-periods of significant differences in the IC time-courses between go and no-go trials are listed in Table 3 and their temporal order is depicted in Figure 4 (upper left panel). The earliest differences occurred at around 70 ms after stimulus presentation in IC1, indicating that the go and no-go stimuli were processed differently already at the early sensory stage. Early differences, starting at around 80 ms, were also observed in the attentional control processes captured by IC4 and IC5. Note that although IC5 seemed to capture the N2/P3 component in the standard ERP, the earliest differences between go and no-go trials seem to reflect smaller yet relevant amplitude fluctuations before the actual development of the N2/P3 wave. Next, significant differences between go and no-go trials occurred in the motor control components (IC2 and IC3) starting from around 130 ms onwards. These were the only components showing hand-specific activity (captured by the factor SIDE), as well as SIDE-by-TRIAL interactions, with the strongest activity in go right and go left trials for IC2 and IC3, respectively. Lastly, there was a short but significant difference between go and no-go trials in the right-lateralized IC6, starting at around 170 ms. No significant differences occurred in the time-course of IC7. In sum, the component activity between go and nogo trials before the no-go prEMG peak latency at around 320 ms revealed significant differences first in terms of sensory processing (IC1), then in attentional control networks (IC4 and IC5), then in motor control networks (IC2 and IC3), and finally in the right-lateralized control network (IC6).

**Figure 4.**
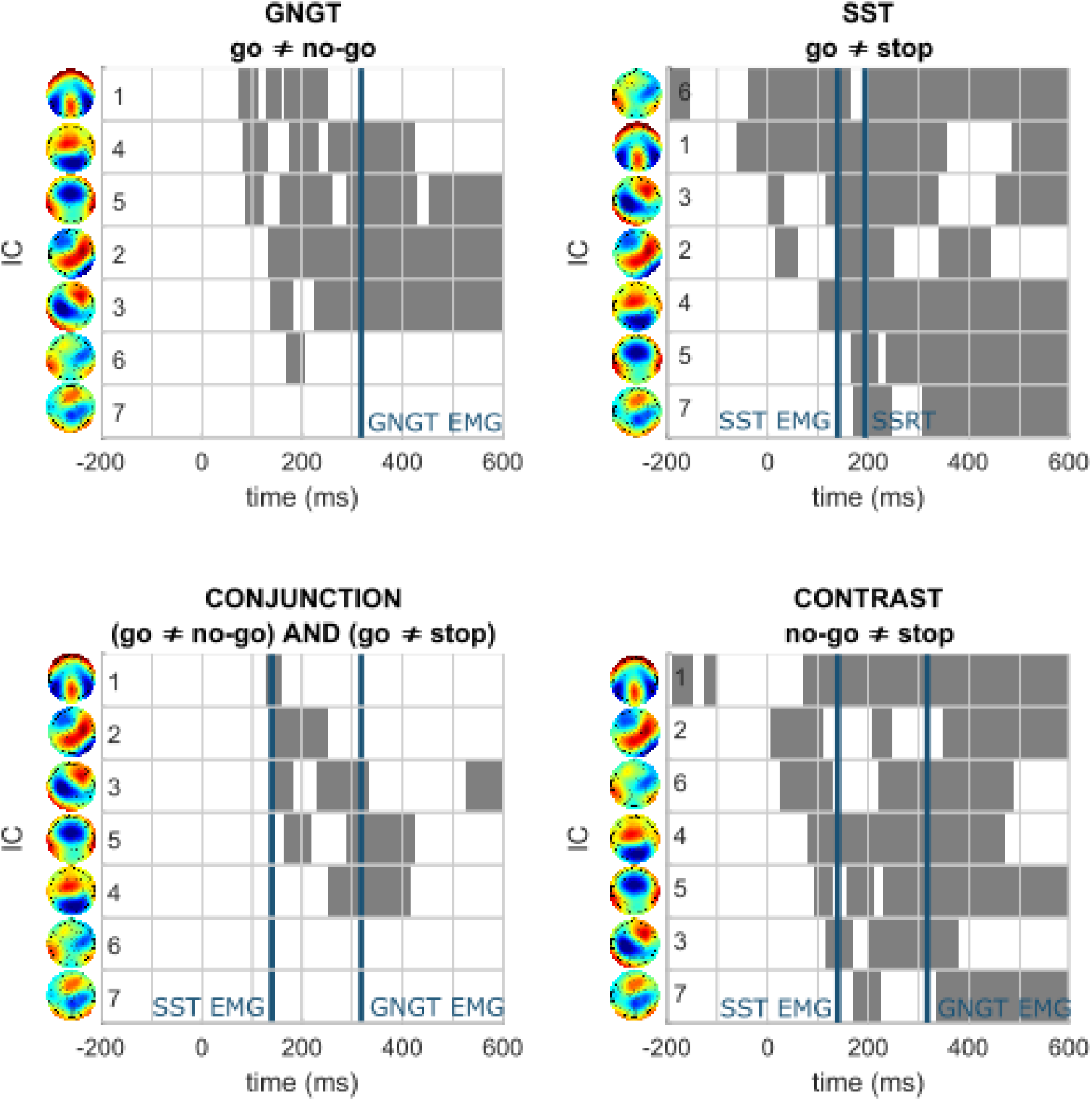
The timing of significant effects in the GNGT (upper left), SST (upper right), the conjunction (lower left), and contrast analysis (lower right). The gray areas mark the significant time intervals for a given effect. ICs are ordered top-to-bottom according to when the significant statistical effect on that IC first occurred. The blue vertical lines represent the prEMG peak latencies or SSRT, as specified on the figure.

*SST*. Time-periods of significant differences between go and stop trials in the SST are listed in Table 3 and in Figure 4 (upper right panel). Differences between go and stop trials occurred already during the pre-stimulus baseline period in the right-lateralized control network IC6, as well as in the early sensory IC1. Very early differences, occurring almost immediately after stimulus presentation, were also observed in the motor control networks IC2 and IC3. These were followed by significant differences in the fronto-parietal IC4 at 96ms. These were the only differences observed before the prEMG peak latency at around 140 ms. Later differences occurred in IC5 that captured the N2/P3 complex, and IC7 from 167 and 170 ms onwards, respectively. In sum, response cancellation in the SST was characterized by pre-stimulus activity in the right-lateralized control and sensory networks, followed by rapid changes in the wider motor control system.

It may be argued that the early differences seen in IC6, IC1, IC2 and IC3 may merely be driven by the processing of the preceding go stimulus in stop trials. However, a number of steps were taken to avoid such contamination during data collection and preprocessing: first, the jittering of the SSD typically attenuates go-related activity as the stop-locked EEG is not time-locked to the go stimulus or RTs; second, go and stop processes are assumed to be independent and would therefore be separable through G-ICA; third, we further separated the overlapping go and stop processes by means of regression. However, the comparisons between go and stop trials suffer from the fact that, according to the horse race model, only slow go responses get inhibited, as fast go responses escape the inhibition process. The observed effects in the pre-stimulus baseline could therefore reflect differences in motor preparation and/or proactive inhibition between fast and slow go responses. We therefore conducted two follow-up analyses: we first contrasted fast and slow go responses, and then successful and unsuccessful stop trials. The results are listed in Table 4 and are visualized in Figure 5. For the sensory IC1, differences occurred between fast and slow go responses at 138-190ms and then again after 370 ms. These differences were reflected also in the contrast of successful and unsuccessful stop trials, starting as early as 8 ms after stop stimulus presentation and continuing throughout the trial. Similarly, the motor control components IC2 and IC3 showed differences between slow and fast responses prior to response execution (here, hand-specificity occurred in the SST as well). Early differences in these ICs occurred also between successful and unsuccessful stop trials. For the right-lateralized IC6, differences occurred between slow and fast go trials prior to response execution, but no such differences were observed between successful and unsuccessful stop trials. In sum, our analyses suggest that the pre-stimulus differences between go and stop trials may be driven by changes between fast and slow go trials, that in turn play a role in whether the ongoing response can be inhibited or not. Altogether, this suggests that at least part of the SST performance and neurophysiological changes appear to reflect earlier modulation of sensory and motor processes by the task context, independent of stop stimulus processing.

**Figure 5.**
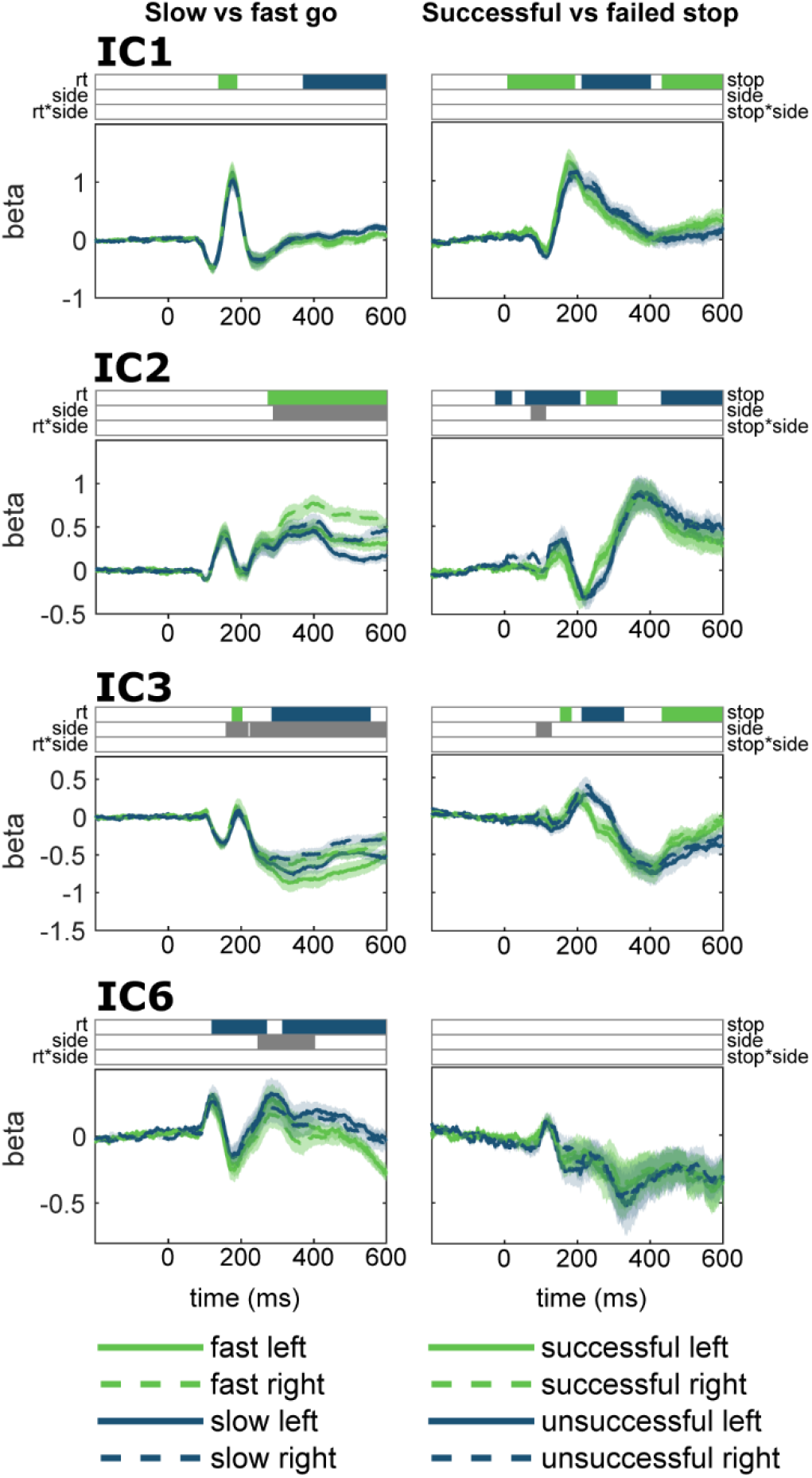
IC time-courses in the SST, comparing slow and fast go trials (left panels), as well as successful and unsuccessful stops (right panel). The bars above the time-courses indicate significant time-intervals for each effect. Green indicates the time-intervals where fast > slow go or successful > unsuccessful stop. Blue indicates time-intervals where slow > fast go or unsuccessful > successful stop.

**Table 4.** The results of the permutation tests comparing slow and fast go responses as well as successful and failed stop trials. The permutation tests indicate the time-periods (ms) of statistical differences of the model RT (slow/fast) + SIDE (left/right) + RT*SIDE and STOP (successful/failed) + SIDE (left/right) + STOP*SIDE. As no interaction effects were significant in either model, these columns are omitted from the table. Alphafdr = 0.05.

| IC | Slow vs fast go |  | Successful vs unsuccessful stop |  |
| --- | --- | --- | --- | --- |
|  | RT | side | stop | side |
| 2 | 138 – 190 | - | 8 – 194 | - |
|  | 370 – 598 |  | 212 – 402 |  |
|  |  |  | 432 – 598 |  |
| 4 | 120 – 272 | 246 – 404 | - | - |
|  | 314 – 598 |  |  |  |
| 6 | 272 – 598 | 286 – 598 | -26 – 20 | 72-114 |
|  |  |  | 56 – 208 |  |
|  |  |  | 224 – 310 |  |
|  |  |  | 430 – 598 |  |
| 8 | 174 – 204 | 158 – 220 | 152 – 184 | 76 – 130 |
|  | 284 – 556 | 224 – 598 | 212 – 328 |  |
|  |  |  | 432 – 598 |  |

#### 3.3.3 Overlapping IC dynamics (go vs. no-go *and* go vs. stop)

The results of the conjunction analysis are listed in Table 3 and Figure 4 (lower left panel). Here, significance indicates those time-periods when no-go vs. go activity was of similar difference as the stop vs. go activity. The earliest conjunction occurred in sensory IC1 at 128 ms, followed by the motor control components IC2 and IC3 at 136 and 142 ms, respectively. This clearly showed that, apart from the similarities in the sensory processing between the two tasks, the earliest similarities occurred at or after the time-point when inhibition processes had already reached the peripheral level in the SST, as shown by the prEMG peak latency. Further conjunction periods occurred in IC5 (N2/P3) starting from 166 ms and in IC4 (fronto-parietal attention) from 250 ms onwards. These are well in line with previous research identifying the N2/P3 complex in both tasks, but their temporal conjunction that appeared later in task processing further supports that they reflect other processes than inhibition, e.g. conflict monitoring, attentional capture or feedback processes. There was no conjunction in the right-lateralized cognitive control component IC6 nor in the IC7.

#### 3.3.4 Contrasting IC dynamics (no-go vs. stop)

There were more differences in the time-courses of no-go and stop trials than there were similarities between the two tasks (Table 3, Figure 4 lower right panel). Converging with the results of the SST analysis, early differences between no-go and stop trials occurred in the right-lateralized control component IC6, then sensory-attentional IC1, and the motor control components IC2 and IC3. Early differences, at around 100-200 ms, were also observed in the attention and cognitive control components IC4, IC5 and IC7. Thus, early differences that continued throughout the trial time-course occurred in all identified components.

## 4. Discussion

Our primary goal was to delineate and compare the cascade of cognitive processes in the GNGT and the SST. We hypothesized that the GNGT and SST may share similar task processing, including similar inhibitory mechanisms. Alternatively, GNGT and SST may operate through different inhibitory mechanisms. We found evidence against the first and in support of the second alternative, suggesting that response inhibition in the most commonly used tasks, the GNGT and SST, rely on different neural mechanisms. This conclusion is based on: 1) behavioral results where go responses were greatly delayed in the SST compared to the GNGT; 2) prEMG activity that declined considerably faster in the SST than in the GNGT; 3) EEG results where the temporal dynamics of ICs differed greatly between the two tasks and showed only very little overlap in the conjunction analysis. The cascade of EEG changes between the go and no-go trials in the GNGT followed an expected and intuitive order, starting from the sensory component, followed by the attentional allocation and/or cognitive control components and only then the motor control components. In the SST, the brain activity was different, suggesting an important role of the cognitive control already at the prestimulus period that might bias the sensory and motor processes prior to the stop stimulus presentation. This might trigger very fast changes in sensory-motor integration upon the presentation of the stop stimulus that are independent of the later more effortful attentional mechanisms.

The first differences between the go and no-go trials in the GNGT occurred in sensory processing, followed by the attentional and the motor control processes. Notably, sensory IC1 was initially stronger in go than in no-go trials, and vice versa in the later time-points. Similarly, there was a strong positive P3-like deflection in go trials both in the ICs (IC4 and 5) and in standard ERPs. EMG results indicate that the electrophysiological effects may be due to a high degree of response competition during action selection in go trials. PrEMG responses were detected more often in the unselected hand in go trials than in the no-go trials, and these declined earlier (289 vs 316 ms for go and no-go trials, respectively). It is possible then that no-go trials recruit mechanisms akin to response selection. In fact, it may be speculated that there is no additional need for an active inhibition mechanism in no-go trials, as all pre-potent but irrelevant responses may be resolved at the response selection stage, similarly to the response competition as seen in the go trials. The late and rather infrequent prEMG observed in the no-go trials may, in this case, reflect trials that escaped the earlier response selection stage. Such an interpretation is in line with the action restraint model, which states that correct no-go performance actually corresponds to the decision not to respond (Bari et al., 2009; Schachar et al., 2007). However, the additional changes in the attentional, cognitive and motor control components, as well as the delayed prEMG peak in the no-go trials may suggest additional processing stages after the initial response selection. Whether these reflect sequential response selection and inhibition processes, or simply confounding cognitive processes elicited by the novelty or surprise, is an important issue for future research.

The SST exhibited differences between the go and stop trials already at the pre-stop level in the right-lateralized control component IC6 and the sensory IC1. Similarly, changes in the motor control components IC2 and IC3 appeared very early after the stop signal (14 and 2 ms, respectively, making it unlikely that these were driven by stop stimulus detection). This early activity appeared to differentiate also between slow and fast go trials, as well as between successful and unsuccessful stop trials (apart from IC6). It is likely then that activity in the sensory and motor networks are modulated by proactive control system in expectation for a stop signal and that the readiness of these systems may influence the performance upon the stop stimulus detection. Remarkably, the actual stopping triggered by the stop-stimulus presentation appears to be very fast, at around 140 ms. This is in line with our findings of peripheral EMG decline at around 150 ms in different versions of SST (Atsma et al., 2018; Raud and Huster, 2017), as well as with studies showing the reduction of corticomotor excitability around this time. Nonetheless, it is earlier than expected from the conceptualization of response inhibition as a higher-order cognitive function (e.g. Miyake et al. 2000; Aron 2007; Diamond 2013).

Converging evidence indicates that response inhibition, as measured in the SST, arises from the interaction of effortful proactive and reflexive reactive mechanisms. For example, if the stop signal is presented for a duration where the participants cannot perceive it consciously, it still results in delayed RTs (van Gaal et al., 2008), yet only if stopping is a possibility in the general task context (Chiu and Aron, 2014). Context-related modulation of motor processes seems to recruit the same right lateralized network that is active in outright stopping (Jahfari et al., 2012), although there seems to be some regional specificity differentiating between the regions of context- and stop-related processes within this network (Messel et. al., submitted). Verbruggen et al. (2014) proposed that stopping an initiated response in the SST is akin to a *prepared* or *intention-based reflex*, where sensory and motor processes are biased towards fast and efficient stop signal processing. Our results fully support this proposal, showing early processing in the control and sensory networks that is associated with slower RTs and followed by rapid changes in the motor control regions upon the detection of stop signal.

Impaired response inhibition is considered one of the core deficits in several psychiatric disorders, particularly in attention-deficit hyperactivity disorder and obsessive compulsive disorder (Lipszyc and Schachar, 2010). In addition, inhibition tasks belong to the standard test batteries for testing the neurocognitive impairments after acquired brain damage, where these tests contribute to the diagnosis of the rather undifferentiated dysexecutive syndrome (Stuss and Alexander, 2007). Our results add to the converging evidence that the practice of using such tasks interchangeably to measure a deficit in inhibitory control is unwarranted. For example, Krämer et al. (2013) found that patients with lateral prefrontal cortex lesions made more no-go omissions than healthy control participants in the GNGT, but were relatively comparable in their SST performance. Our results further indicate that performance in response inhibition tasks is additionally affected by sensory and motor processes, raising the possibility that impaired task performance may stem from processes different from inhibition.

Our approach for the decomposition of EEG via G-ICA was completely data-driven, yet, the recovered IC activations and source constellations converged on the regions similar to those of earlier studies of response inhibition. Nonetheless, one has to be aware that while larger components (e.g. P3) tend to be recovered similarly between different studies, the exact decomposition is dependent on the specific task constellation. Another caveat is that we cannot differentiate processes that show statistical dependencies and therefore converge into a single IC, while their time-courses might reflect functionally distinct processes. For example, the motor control components IC2 and IC3 may include primary motor and premotor regions, but also other prefrontal areas such as IFC and pre-SMA. Given that these IC’s were prominent in both tasks, it might be that the actual inhibition network is still the same in both tasks, yet it receives input from different functional systems, i.e. attentional allocation and action selection in the GNGT and proactive inhibition system in the SST. Thus, the inter-dependencies of the areas within the regions that contribute to each component, as well as causal dependencies between different functional networks should be the priority of future research to fully understand the cascade of processes that ultimately result in suppressed action.

## 5. Conclusion

Based on the behavioral performance, EMG, and EEG-derived independent component activity, we conclude that the most commonly used response inhibition tasks, the GNGT and SST, recruit different brain mechanisms with different temporal dynamics. In both tasks, there is a contribution of early signal detection to successful suppression of a response. In the GNGT, there is an early activation of the fronto-parietal attentional system and only then in the motor control components, resulting in a rather late inhibition at the peripheral level at around 316 ms. In the SST, there seems to be a significant contribution of the frontal control components already prior to the stimulus detection steps and very early changes in the motor control components. This is paralleled by delayed go RTs and fast stopping as indexed via prEMG at around 140 ms. Thus, inhibition in the SST is achieved by proactive biasing of the sensory-motor system in preparation for the stop signal, followed by a fast reflexive inhibitory process within the motor system after the detection of the stop stimulus. Our results do not agree with the notion of a unitary construct of response inhibition and implies that subtle changes in the task instructions can lead to marked changes in the underlying neuro-cognitive processes. This implies caution when comparing different patient populations, as deficits in behavioral performance may be misleadingly assigned to deficient response inhibition.

## Funding

This research was supported by basic funds provided by the Department of Psychology, University of Oslo.

## Abbreviations

GNGT: go/no-go task
G-ICA: group independent component analysis
IC: independent component
prEMG: partial response electromyography
rERP: regression event related potential
SSD: stop signal delay
SSRT: stop signal reaction time
SST: stop signal task

## References

Albares, M., Lio, G., Criaud, M., Anton, J.-L., Desmurget, M., Boulinguez, P., 2014. The dorsal medial frontal cortex mediates automatic motor inhibition in uncertain contexts: evidence from combined fMRI and EEG studies. Hum. Brain Mapp. 35, 5517–5531. https://doi.org/10.1002/hbm.22567

Aron, A.R., 2007. The neural basis of inhibition in cognitive control. Neurosci. Rev. J. Bringing Neurobiol. Neurol. Psychiatry 13, 214–228. https://doi.org/10.1177/1073858407299288

Aron, A.R., Robbins, T.W., Poldrack, R.A., 2014. Inhibition and the right inferior frontal cortex: one decade on. Trends Cogn. Sci. 18, 177–185. https://doi.org/10.1016/j.tics.2013.12.003

Aron, A.R., Robbins, T.W., Poldrack, R.A., 2004. Inhibition and the right inferior frontal cortex. Trends Cogn. Sci. 8, 170–177. https://doi.org/10.1016/j.tics.2004.02.010

Atsma, J., Maij, F., Gu, C., Medendorp, W.P., Corneil, B.D., 2018. Active Braking of Whole-Arm Reaching Movements Provides Single-Trial Neuromuscular Measures of Movement Cancellation. J. Neurosci. Off. J. Soc. Neurosci. 38, 4367–4382. https://doi.org/10.1523/JNEUROSCI.1745-17.2018

Band, G.P.H., van der Molen, M.W., Logan, G.D., 2003. Horse-race model simulations of the stop-signal procedure. Acta Psychol. (Amst.) 112, 105–142.

Bari, A., Eagle, D.M., Mar, A.C., Robinson, E.S.J., Robbins, T.W., 2009. Dissociable effects of noradrenaline, dopamine, and serotonin uptake blockade on stop task performance in rats. Psychopharmacology (Berl.) 205, 273–283. https://doi.org/10.1007/s00213-009-1537-0

Burle, B., Possamaï, C.-A., Vidal, F., Bonnet, M., Hasbroucq, T., 2002. Executive control in the Simon effect: an electromyographic and distributional analysis. Psychol. Res. 66, 324–336. https://doi.org/10.1007/s00426-002-0105-6

Cai, W., Ryali, S., Chen, T., Li, C.-S.R., Menon, V., 2014. Dissociable roles of right inferior frontal cortex and anterior insula in inhibitory control: evidence from intrinsic and task-related functional parcellation, connectivity, and response profile analyses across multiple datasets. J. Neurosci. Off. J. Soc. Neurosci. 34, 14652–14667. https://doi.org/10.1523/JNEUROSCI.3048-14.2014

Chiu, Y.-C., Aron, A.R., 2014. Unconsciously triggered response inhibition requires an executive setting. J. Exp. Psychol. Gen. 143, 56–61. https://doi.org/10.1037/a0031497

Coxon, J.P., Stinear, C.M., Byblow, W.D., 2006. Intracortical inhibition during volitional inhibition of prepared action. J. Neurophysiol. 95, 3371–3383. https://doi.org/10.1152/jn.01334.2005

Cunillera, T., Brignani, D., Cucurell, D., Fuentemilla, L., Miniussi, C., 2016. The right inferior frontal cortex in response inhibition: A tDCS-ERP co-registration study. NeuroImage 140, 66–75. https://doi.org/10.1016/j.neuroimage.2015.11.044

Dambacher, F., Sack, A.T., Lobbestael, J., Arntz, A., Brugman, S., Schuhmann, T., 2014. A network approach to response inhibition: dissociating functional connectivity of neural components involved in action restraint and action cancellation. Eur. J. Neurosci. 39, 821–831. https://doi.org/10.1111/ejn.12425

De Jong, R., Coles, M.G., Logan, G.D., Gratton, G., 1990. In search of the point of no return: the control of response processes. J. Exp. Psychol. Hum. Percept. Perform. 16, 164–182.

Delorme, A., Makeig, S., 2004. EEGLAB: an open source toolbox for analysis of single-trial EEG dynamics including independent component analysis. J. Neurosci. Methods 134, 9–21. https://doi.org/10.1016/j.jneumeth.2003.10.009

Diamond, A., 2013. Executive functions. Annu. Rev. Psychol. 64, 135–168. https://doi.org/10.1146/annurev-psych-113011-143750

Eagle, D.M., Bari, A., Robbins, T.W., 2008. The neuropsychopharmacology of action inhibition: crossspecies translation of the stop-signal and go/no-go tasks. Psychopharmacology (Berl.) 199, 439–456. https://doi.org/10.1007/s00213-008-1127-6

Eichele, T., Rachakonda, S., Brakedal, B., Eikeland, R., Calhoun, V.D., 2011. EEGIFT: Group Independent Component Analysis for Event-Related EEG Data. Comput. Intell. Neurosci. 2011. https://doi.org/10.1155/2011/129365

Enriquez-Geppert, S., Konrad, C., Pantev, C., Huster, R.J., 2010. Conflict and inhibition differentially affect the N200/P300 complex in a combined go/nogo and stop-signal task. NeuroImage 51, 877–887. https://doi.org/10.1016/j.neuroimage.2010.02.043

Filipović, S.R., Jahanshahi, M., Rothwell, J.C., 2000. Cortical potentials related to the nogo decision. Exp. Brain Res. 132, 411–415.

Fujiyama, H., Tandonnet, C., Summers, J.J., 2011. Age-related differences in corticospinal excitability during a Go/NoGo task. Psychophysiology 48, 1448–1455. https://doi.org/10.1111/j.1469-8986.2011.01201.x

Gramfort, A., Papadopoulo, T., Olivi, E., Clerc, M., 2010. OpenMEEG: opensource software for quasistatic bioelectromagnetics. Biomed. Eng. OnLine 9, 45. https://doi.org/10.1186/1475-925X-9-45

Guo, Y., Schmitz, T.W., Mur, M., Ferreira, C.S., Anderson, M.C., 2018. A supramodal role of the basal ganglia in memory and motor inhibition: Meta-analytic evidence. Neuropsychologia 108, 117–134. https://doi.org/10.1016/j.neuropsychologia.2017.11.033

Hasbroucq, T., Possamaï, C.A., Bonnet, M., Vidal, F., 1999. Effect of the irrelevant location of the response signal on choice reaction time: an electromyographic study in humans. Psychophysiology 36, 522–526.

Himberg, J., Hyvarinen, A., 2003. Icasso: software for investigating the reliability of ICA estimates by clustering and visualization, in: 2003 IEEE XIII Workshop on Neural Networks for Signal Processing (IEEE Cat. No.03TH8718). Presented at the 2003 IEEE XIII Workshop on Neural Networks for Signal Processing (IEEE Cat. No.03TH8718), pp. 259–268. https://doi.org/10.1109/NNSP.2003.1318025

Hoshiyama, M., Kakigi, R., Koyama, S., Takeshima, Y., Watanabe, S., Shimojo, M., 1997. Temporal changes of pyramidal tract activities after decision of movement: a study using transcranial magnetic stimulation of the motor cortex in humans. Electroencephalogr. Clin. Neurophysiol. 105, 255–261.

Hoshiyama, M., Koyama, S., Kitamura, Y., Shimojo, M., Watanabe, S., Kakigi, R., 1996. Effects of judgement process on motor evoked potentials in Go/No-go hand movement task. Neurosci. Res. 24, 427–430.

Huster, R.J., Enriquez-Geppert, S., Lavallee, C.F., Falkenstein, M., Herrmann, C.S., 2013. Electroencephalography of response inhibition tasks: functional networks and cognitive contributions. Int. J. Psychophysiol. Off. J. Int. Organ. Psychophysiol. 87, 217–233. https://doi.org/10.1016/j.ijpsycho.2012.08.001

Huster, R.J., Messel, M.S., Thunberg, C., Raud, L., 2019. The P300 as marker of inhibitory control – fact or fiction? bioRxiv 694216. https://doi.org/10.1101/694216

Huster, R.J., Plis, S.M., Calhoun, V.D., 2015. Group-level component analyses of EEG: validation and evaluation. Front. Neurosci. 9, 254. https://doi.org/10.3389/fnins.2015.00254

Huster, R.J., Plis, S.M., Lavallee, C.F., Calhoun, V.D., Herrmann, C.S., 2014. Functional and effective connectivity of stopping. NeuroImage 94, 120–128. https://doi.org/10.1016/j.neuroimage.2014.02.034

Huster, R.J., Raud, L., 2018. A Tutorial Review on Multi-subject Decomposition of EEG. Brain Topogr. 31, 3–16. https://doi.org/10.1007/s10548-017-0603-x

Huster, R.J., Schneider, S., Lavallee, C.F., Enriquez-Geppert, S., Herrmann, C.S., 2017. Filling the void-enriching the feature space of successful stopping. Hum. Brain Mapp. 38, 1333–1346. https://doi.org/10.1002/hbm.23457

Jahfari, S., Verbruggen, F., Frank, M.J., Waldorp, L.J., Colzato, L., Ridderinkhof, K.R., Forstmann, B.U., 2012. How preparation changes the need for top-down control of the basal ganglia when inhibiting premature actions. J. Neurosci. Off. J. Soc. Neurosci. 32, 10870–10878. https://doi.org/10.1523/JNEUROSCI.0902-12.2012

Johnstone, S.J., Dimoska, A., Smith, J.L., Barry, R.J., Pleffer, C.B., Chiswick, D., Clarke, A.R., 2007. The development of stop-signal and Go/Nogo response inhibition in children aged 7-12 years: performance and event-related potential indices. Int. J. Psychophysiol. Off. J. Int. Organ. Psychophysiol. 63, 25–38. https://doi.org/10.1016/j.ijpsycho.2006.07.001

Krämer, U.M., Solbakk, A.-K., Funderud, I., Løvstad, M., Endestad, T., Knight, R.T., 2013. The role of the lateral prefrontal cortex in inhibitory motor control. Cortex J. Devoted Study Nerv. Syst. Behav. 49, 837–849. https://doi.org/10.1016/j.cortex.2012.05.003

Kybic, J., Clerc, M., Abboud, T., Faugeras, O., Keriven, R., Papadopoulo, T., 2005. A common formalism for the Integral formulations of the forward EEG problem. IEEE Trans. Med. Imaging 24, 12–28. https://doi.org/10.1109/TMI.2004.837363

Langford, Z.D., Krebs, R.M., Talsma, D., Woldorff, M.G., Boehler, C.N., 2016a. Strategic downregulation of attentional resources as a mechanism of proactive response inhibition. Eur. J. Neurosci. https://doi.org/10.1111/ejn.13303

Langford, Z.D., Schevernels, H., Boehler, C.N., 2016b. Motivational context for response inhibition influences proactive involvement of attention. Sci. Rep. 6, 35122. https://doi.org/10.1038/srep35122

Liebrand, M., Kristek, J., Tzvi, E., Krämer, U.M., 2018. Ready for change: Oscillatory mechanisms of proactive motor control. PloS One 13, e0196855. https://doi.org/10.1371/journal.pone.0196855

Lipszyc, J., Schachar, R., 2010. Inhibitory control and psychopathology: a meta-analysis of studies using the stop signal task. J. Int. Neuropsychol. Soc. JINS 16, 1064–1076. https://doi.org/10.1017/S1355617710000895

Littman, R., Takács, Á., 2017. Do all inhibitions act alike? A study of go/no-go and stop-signal paradigms. PloS One 12, e0186774. https://doi.org/10.1371/journal.pone.0186774

Logan, G.D., Cowan, W.B., 1984. On the ability to inhibit thought and action: A theory of an act of control. Psychol. Rev. 91, 295–327. https://doi.org/10.1037/0033-295X.91.3.295

Macdonald, H.J., Coxon, J.P., Stinear, C.M., Byblow, W.D., 2014. The Fall and Rise of Corticomotor Excitability with Cancellation and Reinitiation of Prepared Action. J. Neurophysiol. https://doi.org/10.1152/jn.00366.2014

McNab, F., Leroux, G., Strand, F., Thorell, L., Bergman, S., Klingberg, T., 2008. Common and unique components of inhibition and working memory: an fMRI, within-subjects investigation. Neuropsychologia 46, 2668–2682. https://doi.org/10.1016/j.neuropsychologia.2008.04.023

Messel, M.S., Raud, L., Hoff, P.K., Skaftnes, C.S., Huster, R.J., submitted. Strategy switched in proactive inhibitory control and their association with task-general and stopping-specific networks.

Miyake, A., Friedman, N.P., Emerson, M.J., Witzki, A.H., Howerter, A., Wager, T.D., 2000. The unity and diversity of executive functions and their contributions to complex “Frontal Lobe” tasks: a latent variable analysis. Cognit. Psychol. 41, 49–100. https://doi.org/10.1006/cogp.1999.0734

Nee, D.E., Wager, T.D., Jonides, J., 2007. Interference resolution: insights from a meta-analysis of neuroimaging tasks. Cogn. Affect. Behav. Neurosci. 7, 1–17.

Nichols, T., Brett, M., Andersson, J., Wager, T., Poline, J.-B., 2005. Valid conjunction inference with the minimum statistic. NeuroImage 25, 653–660. https://doi.org/10.1016/j.neuroimage.2004.12.005

Raud, L., Huster, R.J., 2017. The Temporal Dynamics of Response Inhibition and their Modulation by Cognitive Control. Brain Topogr. 30, 486–501. https://doi.org/10.1007/s10548-017-0566-y

Rochet, N., Spieser, L., Casini, L., Hasbroucq, T., Burle, B., 2014. Detecting and correcting partial errors: Evidence for efficient control without conscious access. Cogn. Affect. Behav. Neurosci. 14, 970–982. https://doi.org/10.3758/s13415-013-0232-0

Rubia, K., Russell, T., Overmeyer, S., Brammer, M.J., Bullmore, E.T., Sharma, T., Simmons, A., Williams, S.C., Giampietro, V., Andrew, C.M., Taylor, E., 2001. Mapping motor inhibition: conjunctive brain activations across different versions of go/no-go and stop tasks. NeuroImage 13, 250–261. https://doi.org/10.1006/nimg.2000.0685

Schachar, R., Logan, G.D., Robaey, P., Chen, S., Ickowicz, A., Barr, C., 2007. Restraint and cancellation: multiple inhibition deficits in attention deficit hyperactivity disorder. J. Abnorm. Child Psychol. 35, 229–238. https://doi.org/10.1007/s10802-006-9075-2

Sebastian, A., Pohl, M.F., Klöppel, S., Feige, B., Lange, T., Stahl, C., Voss, A., Klauer, K.C., Lieb, K., Tüscher, O., 2013. Disentangling common and specific neural subprocesses of response inhibition. NeuroImage 64, 601–615. https://doi.org/10.1016/j.neuroimage.2012.09.020

Skippen, P., Fulham, W.R., Michie, P.T., Matzke, D., Heathcote, A., Karayanidis, F., 2019. Reconsidering electrophysiological markers of response inhibition in light of trigger failures in the stop-signal task. bioRxiv 658336. https://doi.org/10.1101/658336

Smith, N.J., Kutas, M., 2015. Regression-based estimation of ERP waveforms: I. The rERP framework. Psychophysiology 52, 157–168. https://doi.org/10.1111/psyp.12317

Smith, S.M., Nichols, T.E., 2009. Threshold-free cluster enhancement: addressing problems of smoothing, threshold dependence and localisation in cluster inference. NeuroImage 44, 83–98. https://doi.org/10.1016/j.neuroimage.2008.03.061

Stuss, D.T., Alexander, M.P., 2007. Is there a dysexecutive syndrome? Philos. Trans. R. Soc. Lond. B. Biol. Sci. 362, 901–915. https://doi.org/10.1098/rstb.2007.2096

Swick, D., Ashley, V., Turken, U., 2011. Are the neural correlates of stopping and not going identical? Quantitative meta-analysis of two response inhibition tasks. NeuroImage 56, 1655–1665. https://doi.org/10.1016/j.neuroimage.2011.02.070

Tadel, F., Baillet, S., Mosher, J.C., Pantazis, D., Leahy, R.M., 2011. Brainstorm: A User-Friendly Application for MEG/EEG Analysis [WWW Document]. Comput. Intell. Neurosci. https://doi.org/10.1155/2011/879716

van Campen, A.D., Neubert, F.-X., van den Wildenberg, W.P.M., Ridderinkhof, K.R., Mars, R.B., 2013. Paired-pulse transcranial magnetic stimulation reveals probability-dependent changes in functional connectivity between right inferior frontal cortex and primary motor cortex during go/no-go performance. Front. Hum. Neurosci. 7, 736. https://doi.org/10.3389/fnhum.2013.00736

van den Wildenberg, W.P.M., Burle, B., Vidal, F., van der Molen, M.W., Ridderinkhof, K.R., Hasbroucq, T., 2010. Mechanisms and dynamics of cortical motor inhibition in the stop-signal paradigm: a TMS study. J. Cogn. Neurosci. 22, 225–239. https://doi.org/10.1162/jocn.2009.21248

van Gaal, S., Ridderinkhof, K.R., Fahrenfort, J.J., Scholte, H.S., Lamme, V.A.F., 2008. Frontal cortex mediates unconsciously triggered inhibitory control. J. Neurosci. Off. J. Soc. Neurosci. 28, 8053–8062. https://doi.org/10.1523/JNEUROSCI.1278-08.2008

Verbruggen, F., McLaren, I.P.L., Chambers, C.D., 2014. Banishing the Control Homunculi in Studies of Action Control and Behavior Change. Perspect. Psychol. Sci. J. Assoc. Psychol. Sci. 9, 497–524. https://doi.org/10.1177/1745691614526414

Wessel, J.R., 2017. Prepotent motor activity and inhibitory control demands in different variants of the go/no-go paradigm. Psychophysiology. https://doi.org/10.1111/psyp.12871

Wessel, J.R., Aron, A.R., 2015. It’s not too late: the onset of the frontocentral P3 indexes successful response inhibition in the stop-signal paradigm. Psychophysiology 52, 472–480. https://doi.org/10.1111/psyp.12374

Winkler, A.M., Ridgway, G.R., Webster, M.A., Smith, S.M., Nichols, T.E., 2014. Permutation inference for the general linear model. NeuroImage 92, 381–397. https://doi.org/10.1016/j.neuroimage.2014.01.060

Yamanaka, K., Kimura, T., Miyazaki, M., Kawashima, N., Nozaki, D., Nakazawa, K., Yano, H., Yamamoto, Y., 2002. Human cortical activities during Go/NoGo tasks with opposite motor control paradigms. Exp. Brain Res. 142, 301–307. https://doi.org/10.1007/s00221-001-0943-2

Zheng, D., Oka, T., Bokura, H., Yamaguchi, S., 2008. The key locus of common response inhibition network for no-go and stop signals. J. Cogn. Neurosci. 20, 1434–1442. https://doi.org/10.1162/jocn.2008.20100

